# Tissue stiffness mapping by light sheet elastography

**DOI:** 10.1101/2023.12.09.570896

**Authors:** Min Zhu, Kaiwen Zhang, Evan C. Thomas, Ran Xu, Brian Ciruna, Sevan Hopyan, Yu Sun

## Abstract

Tissue stiffness plays a crucial role in regulating morphogenesis. The ability to measure and monitor the dynamic progression of tissue stiffness is important for generating and testing mechanistic hypotheses. Methods to measure tissue properties *in vivo* have been emerging but present challenges with spatial and temporal resolution especially in 3D, by their reliance on highly specialised equipment, and/or due to their invasive nature. Here, introduce light sheet elastography, a noninvasive method that couples low frequency shear waves with light sheet fluorescence microscopy by adapting commercially available instruments. With this method, we achieved *in toto* stiffness mapping of organ-stage mouse and zebrafish embryos at cellular resolution. Versatility of the method enabled time-lapse stiffness mapping during tissue remodelling and of the beating embryonic heart. This method expands the spectrum of tools available to biologists and presents new opportunities for uncovering the mechanical basis of morphogenesis.

## INTRODUCTION

Morphogenesis involves complex coordination of biochemical and biophysical signals. Stiffness, a key mechanical property of tissue, plays a multifaceted role by modulating cell behaviours^1–3^ and interacting with biochemical mediators^4–6^. A wholistic understanding of how the distributions and dynamic changes in tissue stiffness contribute to development requires new measurement tools.

Atomic force microscopy (AFM) indentation is a widely used method for measuring 2D stiffness at the tissue surface^7^. To measure the stiffness of deeper tissue layers, surgical removal of the surface layer is required^2^. The few state-of-the-art untethered methods permitting *in vivo* measurement of 3D tissue stiffness require the injection of magnetic beads or droplets (magnetic tweezers)^3,8–10^ or silica beads (optical tweezers)^11^. These methods are cumbersome, cause potential damage especially to sensitive tissues such as the developing brain and heart, and are limited in spatial resolution that relies on the positions of the deposited beads. Importantly, morphogenesis unfolds over time and invasive techniques fail to capture the temporal dynamics of stiffness, thereby limiting current studies to endpoint analyses. An ideal method would be noninvasive, permit high spatial and temporal resolution, and be sufficiently versatile for a substantial proportion of biologists using commercially available equipment.

Elastography is a noninvasive approach that consists of an actuation component to introduce local tissue deformation and an imaging tool to record the response of the tissue to quantify stiffness^12^. In terms of actuation principles, current elastography techniques can be categorised into optical and mechanical modes. In Brillouin microscopy^13,14^, the frequency shift of incident laser Brillouin scattering (i.e., Brillouin frequency shift in GHz) is used as a proxy for tissue stiffness (i.e., longitudinal modulus) assuming known refractive indices. The relationship between Brillouin microscopy measured longitudinal modulus and elastic modulus (a common metric of tissue stiffness), however, remains unclear. A phenomenological positive correlation between the two quantities^15–17^ and no correlation^18^ were both reported when benchmarked against AFM indentation. This is potentially due to the fact that Brillouin microscopy naturally couples mechanical and optical properties of each location in a tissue^13,18^. In terms of measurement speed, the acquisition speed of Brillouin microscopy is typically 20-500 ms per pixel (i.e., 20-500 s for capturing a 1000 × 1000 image)^14^. To solve the reliance on known refractive indices of different locations in a tissue, Brillouin microscopy has been used together with optical diffraction tomography^19^, which further lengthens its acquisition speed. Moreover, Brillouin frequency shift has been reported to be tenfold more sensitive to water content than to stiffness^20,21^. Interpreting Brillouin microscopy results in tissues with varying hydration levels requires caution^13^. Therefore, despite its capability of offering subcellular resolution, Brillouin microscopy is suboptimal for stiffness mapping of 3D tissue comprising multiple slices.

For mechanically actuated elastography, compression optical coherence elastography (C-OCE) and wave-based elastography have been developed^22^. C-OCE, which is based on local strain imaging under axial (depth) compression ^23–25^ relies on known refractive indices and has reduced accuracy and sensitivity when measuring tissues lacking mechanical stiffness contrast^22,26^. For quantitative stiffness measurement, uniform stress is assumed, which introduces errors in mapping geometrically complex tissues with variable stress^22,25^. C-OCE also has limited axial resolution for strain measurement (∼300 µm) in tissues^25^. In wave-based elastography, the propagation speed of a shear wave is proportional to tissue stiffness^27^. This approach has been applied clinically using ultrasound elastography (UE) and 2D/3D magnetic resonance elastography (MRE) for diagnosing stiffness-related diseases such as liver fibrosis^28,29^. Switching the imaging modality to optical microscopy has improved spatial resolution. However, a high actuation frequency in the kHz range has been required to enhance local phase contrast for generating an observable phase shift to calculate shear wave propagation speed. This requirement arises from limited displacement tracking under brightfield imaging with particle imaging velocimetry^30^. Capturing sample response to such high-frequency actuation makes it incompatible with confocal or light sheet imaging and obviates 3D stiffness mapping. The high frequency actuation also leads to rapid amplitude attenuation, impeding whole embryo penetration.

To tackle these limitations, we developed a method for mapping stiffness by coupling mechanical actuation with light sheet fluorescence microscopy (Supplementary Table). A standard piezoelectric device was used to propagate shear waves through organ-stage mouse embryos (∼3 mm). The shear wave displaced cells that were individually visualised by a transgenic far-red nuclear reporter H2B-miRFP703^31^ that offered high-contrast and deep-tissue imaging. Application of a customised pattern-guided cell motion tracking program enabled accurate reconstruction of local displacement patterns and distinguished phase shifts as low as 0.0023 rad at a low actuation frequency (≤10 Hz). We achieved whole embryo shear wave penetration and *in toto* tissue stiffness mapping at cellular (15 µm) resolution with time-lapse capability. Stiffness of the beating embryonic heart during ventricular contraction and relaxation was also mapped for the first time.

## RESULTS

### Light sheet elastography device

To optimise live imaging for mouse embryos, our device was built on a standard light sheet fluorescence microscope that provides high spatiotemporal resolution and minimises photobleaching. Simultaneous planar laser excitation and acquisition enabled by light sheet microscopy mitigated the confounding delay for specimen scanning (illuminating) by confocal microscopes. The light sheet elastography (LSE) device consisted of a standard piezoelectric actuator (P-840.20, Physik Instrumente) that vibrated with displacement of up to ∼30 µm perpendicular to a capillary tube housing an agarose plug. Piezoelectric actuation by a sinusoidal signal generated a shear wave through the agarose plug and the embryo embedded within it (Methods). A mounting stage was secured to the microscope sample stage by permanent magnets (Figures 1A and 1B).

**Figure 1.**
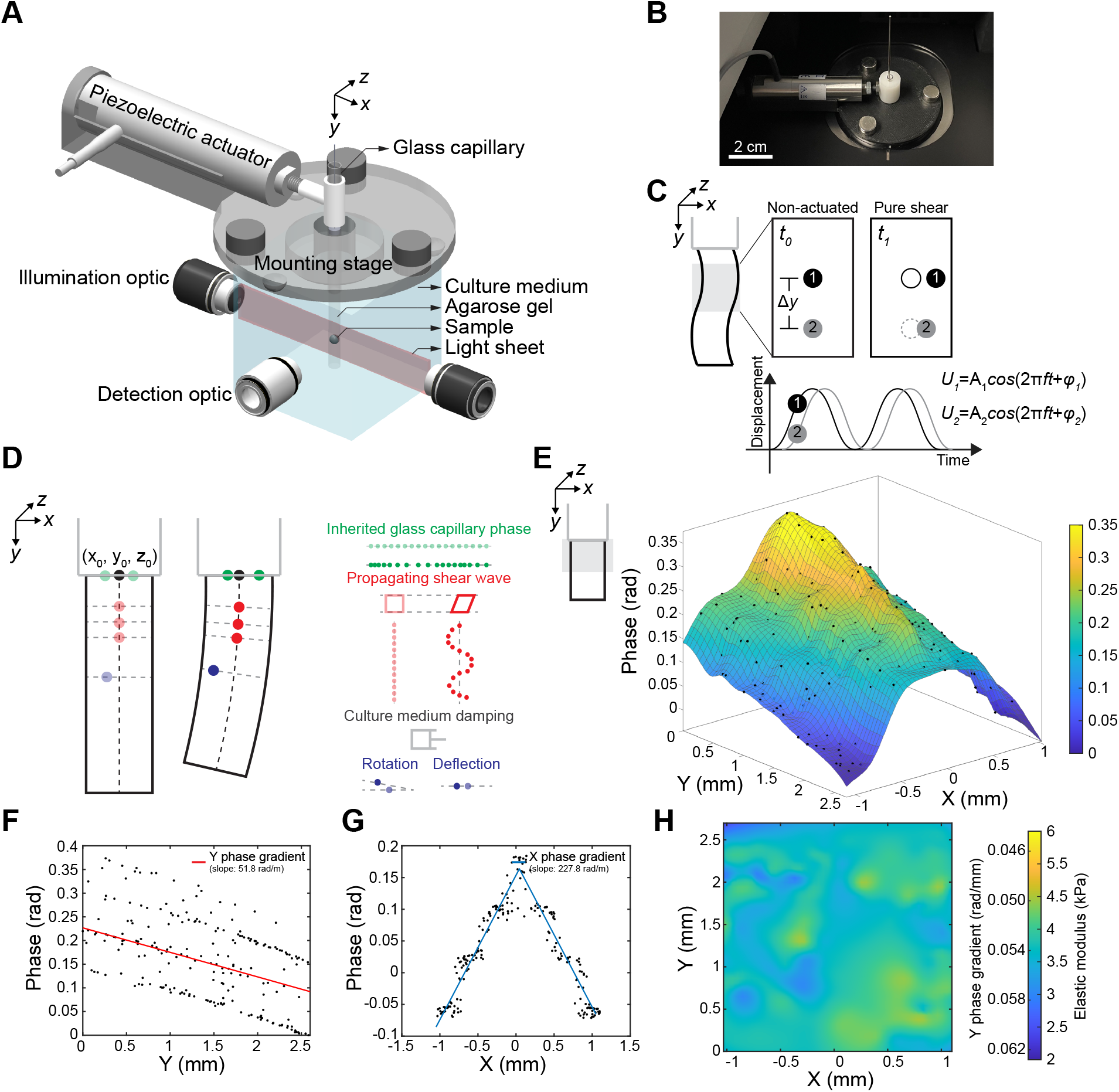
Light sheet elastography. (**A**) 3D structure of the LSE device. (**B**) Experimental setup of LSE. (**C**) Schematic illustrating ideal pure shear wave propagation causing a phase delay along Y axis. (**D**) Schematic describing potential contributors to the local phase. (**E**) 3D rendered phase map of 1% agarose gel under 10 Hz actuation. (**F**) YZ projection of the phase map shown in E showing linear phase decay induced by shear wave propagation along Y axis. (**G**) Inherited glass capillary phase along X axis. (**H**) Stiffness map of 1% agarose gel. Colour-coded scale showing elastic modulus and its corresponding Y phase gradient under 10 Hz actuation.

### Dynamic phase analysis

Under ideal pure shear actuation as illustrated in Figure 1C and Movie S1, the local shear modulus *G* relates to local phase difference by^27^

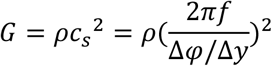

where *ρ* is density; *c*_*s*_ is shear wave propagation speed; *f* is actuation frequency; and Δ*φ*/Δ*y* is the Y phase gradient. The relationship was based on the principle that a forward-propagating wave passes specified point locations with a phase delay. For two points along the path of a shear wave that are in close proximity such that the material in between them can be considered homogeneous, the difference in the local phases at the two points was used to calculate the stiffness of the material between the two points. The local shear modulus *G* is related to elastic modulus *E* (a common metric for describing tissue stiffness) by

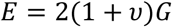

where *υ* is Poisson’s ratio which is typically 0.5 for soft tissue^9,32^.

In addition to the phase arising from pure shear wave propagation, other potential contributors to the phase of the extruded agarose gel are the phase inherited from the glass capillary above the gel, the damping effect of the culture medium, and the geometry (i.e., diameter and extrusion length) of the gel (Figure 1D and Supplementary Note).

To validate the phase-stiffness equation and evaluate the effect of potential contributing factors, we conducted shear wave actuation on 1% agarose gel that contained fluorescent beads. By tracking the displacement of the beads and fitting with the sinusoidal equation to decouple phase values (Methods), we obtained a local phase map of 1% agarose gel (Figure 1E). The phase map decays linearly along Y due to shear wave propagation with an average Y phase gradient of 51.8 rad/m. In addition to the Y phase, the phase also shows a linear symmetric decay along X with a ruffle fold that is inherited from the agarose gel inside the glass capillary (Figure 1G). The decay and ruffle fold resulted from wave propagation in the poroelastic material arising from the glass-gel boundary and wave interference, respectively^33^ (Figure S1A and Supplementary Note). Inside the glass capillary, the agarose gel phase map exhibited the same X phase pattern as the extruded part without Y phase decay (Figures S1B-S1D). This X phase pattern was experimentally determined to be independent of Z location, glass capillary diameter, and agarose gel concentration (Figures S1E-S1G). The X phase speed (i.e., wave propagation speed) remained constant across different actuation frequencies, suggesting that the wave was non-dispersive (Figure S1H).

The extruded gel was found to be critically damped (i.e., damping ratio close to 1) by DMEM (Figure S2A and Supplementary Note). The critical damping effect filters out undesired wave reflection and mitigates its contribution to the local phase under forced oscillation^27^. DMEM damping arose from the added mass of the fluid that was displaced by the agarose gel and viscous fluid shear. From experimental and numerical simulation results, DMEM damping was dominated by mass damping (Figures S2B-S2F). Replacing the culture medium with fluids of different viscosities did not affect the phase result, as they all had a similar density to water. The critical damping effect extended to agarose gels of different elastic moduli and diameters (Figures S2G and S2H). We further verified that the rotational bending and deflection induced by the agarose gel’s geometry was minimal and can be disregarded for phase analysis (Figures S2I-S2L and Supplementary Note). By performing actuation at different frequencies, Y phase speed was found to be constant (i.e., non-dispersive) as depicted in Figure S2M, indicating the predominantly elastic behaviour of the agarose gel. The elastic behaviour of agarose gel was recapitulated by atomic force microscope (AFM) indentation (Figure S3A and Methods).

Hence, the phase in the extruded agarose gel was solved according to

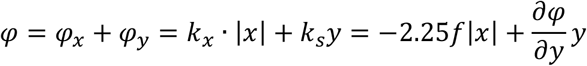

where *k*_*x*_ and *k*_*s*_ are X and Y wavenumbers, respectively (Supplementary Note). Using our customised dynamic phase program (code link in Methods), the partial derivative of phase with respect to Y was computed, and the stiffness of 1% agarose gel was decoupled (Figure 1H) and determined to be 4.1 kPa on average. To validate LSE-measured stiffness, we compared the stiffness of 0.75% to 3% agarose gels quantified by LSE and by AFM indentation (Figures S3B and S3C). LSE-measured stiffness values of agarose gel agreed well with AFM indentation results for gel concentrations ranging from 0.75% to 2% (i.e., <∼10 kPa) with differences less than 7.4%. For 3% agarose gel, LSE-measured values deviated from AFM indentation results by 20.9%. This discrepancy occurred because the frame rate of the Zeiss Z1 light sheet microscope at 1000 × 1000 pixels was unable to capture the rapid propagation of shear waves to decouple the Y phase gradient in the stiff gel. For measuring stiff samples, the LSE pixel setting can be lowered to increase the frame rate. For 3% agarose gel, lowering the pixel setting from 1000 × 1000 to 250 × 250 enabled LSE to reproduce AFM indentation results (within 7.2% difference and reduced variance) at the cost of a smaller field-of-view.

To simulate a heterogenous stiffness environment, we also conducted validation on gradient agarose gels with spatially varying stiffness (Figure S3D and Methods). For 1% to 2% gradient agarose gel, LSE measured continuous points and resolved the stiffness gradient to within 7.7% difference compared to AFM indentation (Figures S3E and S3F).

### *In toto* tissue stiffness mapping

To achieve maximum imaging penetration depth, we performed tissue stiffness mapping of an E9.25 mouse embryo using a transgenic mouse strain harbouring a far-red nuclear reporter H2B-miRFP703^31^. This mouse line has been useful in various live imaging studies as it allows clear visualistion throughout tissue with minimal phototoxicity^3,6,34^. To track the fast moving, densely populated nuclei under shear wave propagation, we developed a motion tracking program utilising a subpixel image registration algorithm^35^ (Figures S4A-S4D, S5A, Movie S2, and code link in Methods), capable of resolving a phase shift as small as 0.0023 rad (Methods). At E9.25, the mouse embryonic heart regularly beats at ∼2 Hz (Figure 2A and Movie S3) with an amplitude that extends to ∼200 µm away from the heart (Figure S5B). To compensate for the heartbeat, we chose our actuation frequency to be distinct from that of the beating frequency and eliminated the heartbeat signal using bandpass filtering (Figure S5A). The time resolution of LSE (i.e., number of frames required for a stable phase reading) was determined to be 7 frames (0.12 s) (Figure S5C). The spatial resolution was calculated as a function of the time resolution, actuation frequency, and sample stiffness considering a minimum signal-to-noise ratio of 3 (Methods). Within the stiffness range of most embryonic tissues (a few hundred Pa), neighbouring cells’ phase difference could be resolved by LSE yielding cellular resolution (<15 µm) for stiffness mapping (Figures S5D, S5E, and Supplementary Note). No significant changes in cell apoptosis were observed after LSE actuation as examined by lysotracker staining (Figure S5F and methods).

**Figure 2.**
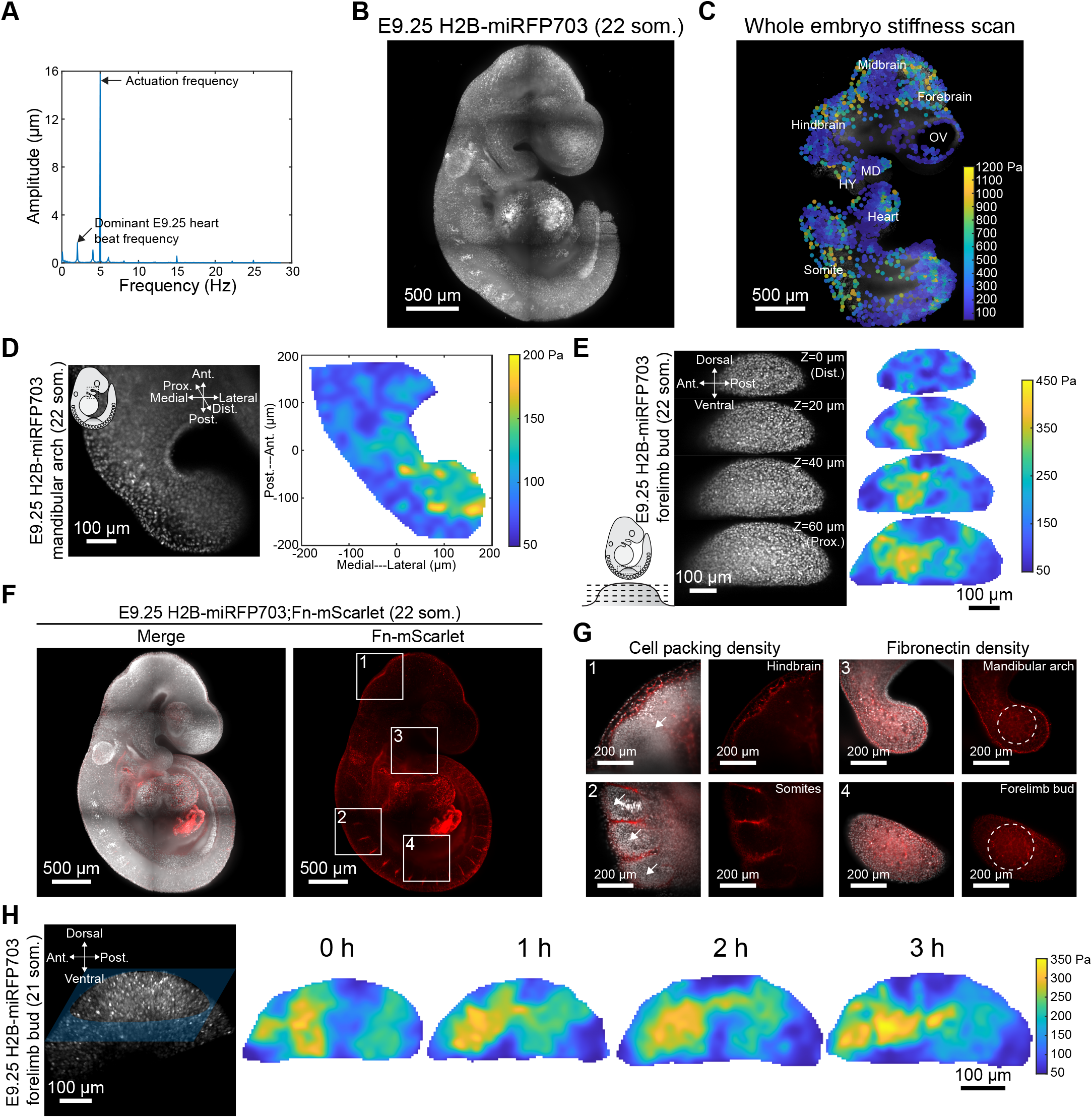
Non-invasive tissue stiffness mapping. (**A**) Fast Fourier transform of a heart cell under 5 Hz actuation. (**B**) 3D rendering of an E9.25 H2B-miRFP703 mouse embryo. (**C**) Slice view of a E9.25 whole embryo stiffness map. In this wide-field depiction of six stitched tiles of 15 μm resolution stiffness data, a subset of data points is shown as individual dots for easy of visualisation. OV, optic vesicle; MD, mandibular arch; HY, hyoid arch. (**D**) Slice view of colour map rendered E9.25 mandibular arch stiffness. (**E**) Slices at different depths of E9.25 limb bud 3D stiffness map. (**F**) 3D rendering of an E9.25 H2B-miRFP703;Fn-mScarlet mouse embryo.(**G**) Contributing factors of tissue stiffness in different regions (solid squares in F). Left panel illustrating high cell packing density (arrows) leads to the stiff hindbrain and somites as shown in (C). Right panel illustrating FN density (dashed circles) modulates stiffness gradient observed in mandibular arch (D) and forelimb bud (E). (**H**) Time-lapse stiffness map of an E9.25 limb bud.

The stiffness of E9.25 mouse embryos was mapped by stitching sagittal plane tiles as shown in Figures 2B and 2C. The void areas in the stiffness map are cavities filled with intraembryonic fluid. Since shear waves do not propagate through fluid^36^, cavities do not affect stiffness mapping results. The E9.25 embryo exhibited a stiff forebrain, hindbrain and somites with comparatively soft midbrain and optic vesicle regions. We compared LSE with established *in vivo* stiffness measurement methods. LSE results agreed well with previously reported values for the E9.25 mandibular arch, mouse forelimb bud, and the tailbud of 16 hpf zebrafish harbouring an mScarlet nuclear reporter^37^ that were measured using AFM indentation^7,38^ and magnetic tweezers^3,8,10^ but with greater spatial detail (Figures 2D, 2E, S5G and S5H).

Cell packing density and extracellular matrix (ECM) composition are regarded as two major contributing factors of tissue stiffness^3,6,39^. Both parameters vary spatially during development. By crossing the nuclear reporter H2B-miRFP703 strain with our endogenous fibronectin (FN, a major ECM protein) reporter strain (Fn-mScarlet)^6^, we explored the potential contributions of these two factors to stiffness (Figure 2F and Methods). Some regions of high stiffness by LSE such as the hindbrain and somites corresponded spatially to high cell packing density whereas others in the mandibular arch and forelimb bud matched FN gradients (Figure 2G). It will be interesting to determine how different determinants of tissue stiffness affect morphogenesis.

### Time-lapse tissue stiffness mapping

The noninvasive nature of LSE enabled time-lapse tissue stiffness mapping. We previously discovered an anteriorly biased tissue stiffness gradient in the mouse E9.25 forelimb bud by taking single time point measurements per embryo^3^. In contrast, LSE enabled time-lapse stiffness mapping over a 3 h period. During that period, the stiffness gradient evolved such that the anterior bias expanded toward the central region (Figure 2H). That progression is consistent with an anterior to central transition of the distal peak of the limb bud as it expands between E9.25-E9.5^3,6^. The ability to observe live changes in tissue stiffness may be useful for defining the basis of morphogenetic movements.

The noninvasive nature of LSE also allowed for stiffness mapping of mechanosensitive tissues such as the beating embryonic heart that cannot be achieved by existing methods. Experiments confirmed that 10 Hz LSE actuation for 1.7 seconds (i.e., 100 frames) did not induce any heartbeat frequency change or arrhythmia (Figure 3A). An important aspect of measuring heart stiffness is that shear wave propagation speed is affected by tissue stress^40^. To differentiate heart stiffness during ventricular relaxation and contraction, we first measured the time weighted average (>10 heartbeats) of E9.25 heart stiffness as shown in Figures 3B and 3C. A time resolution of 0.12 s (i.e., 7 frames) allowed us to isolate stiffness during ventricular relaxation as shown in Figure 3D. Heart stiffness during ventricular contraction was decoupled according to

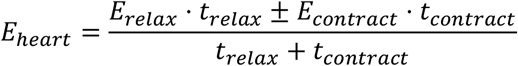

and was 23.2% higher than in the relaxed state (Figures 3E and 3F). Whether the heartbeat increases or decreases the apparent stiffness depends on tissue stress which is determined by the superposition of contractile stress upon initial stress in the relaxed state. An analogy would be stretching (increase) or releasing (decrease) a pre-stressed rubber band. Since LSE can distinguish contributions of contractile stress to stiffness throughout the cardiac cycle, the method can potentially identify the precise functional consequences of deleterious mutations.

**Figure 3.**
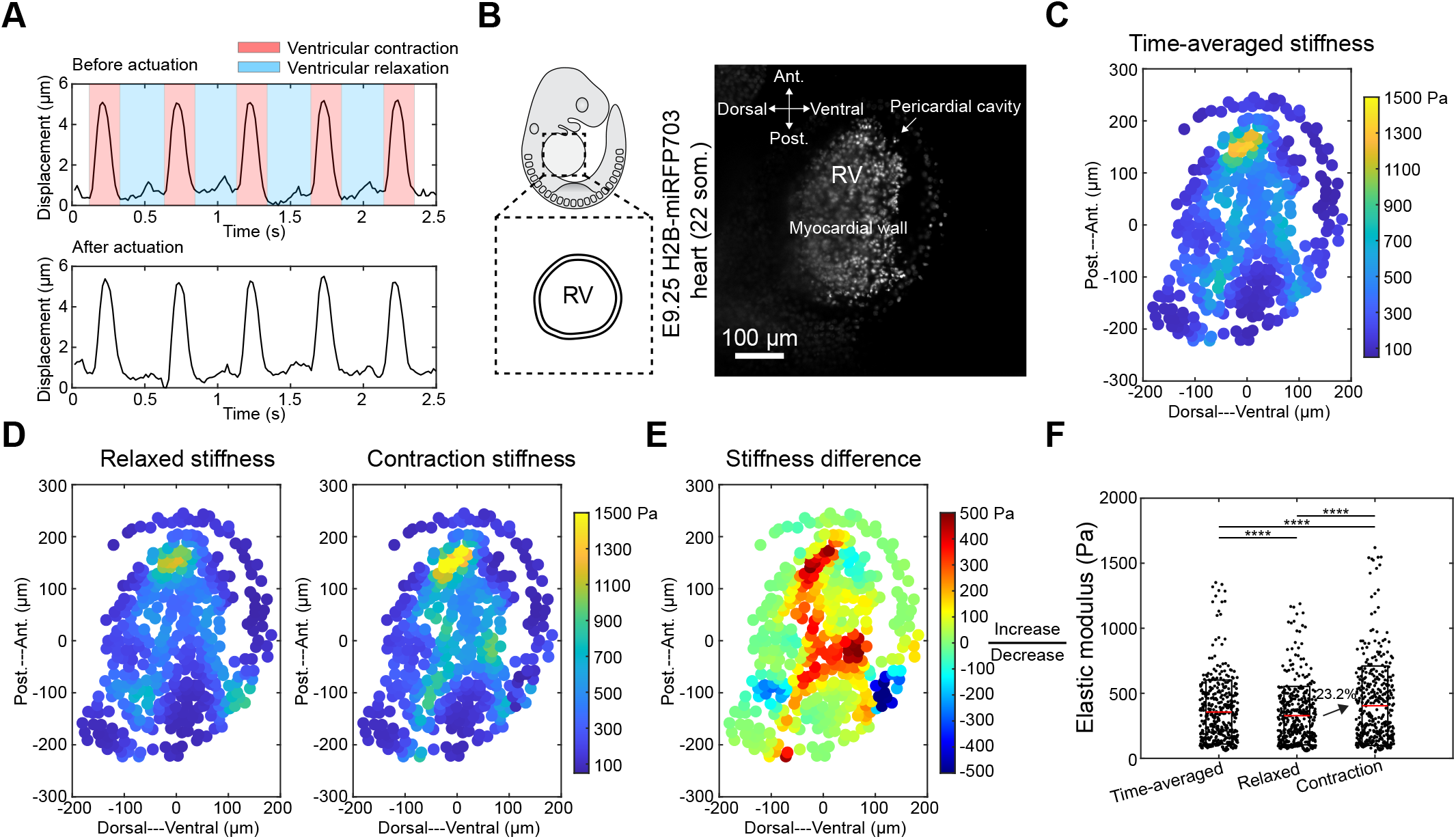
Heart stiffness mapping. (**A**) Heart nuclear displacements by tracking nuclei before and after actuation showing no changes in beating frequency and rhythm. (**B**) Slice view of an E9.25 H2B-miRFP703 embryonic heart showing the focal plane of stiffness mapping. RV, right ventricle. (**C**) Time-averaged heart stiffness map. (**D**) Heart stiffness mapped during ventricular relaxation and decoupled ventricular contraction stiffness map. (**E**) Heart stiffness difference between ventricular relaxation and contraction. (**F**) Box plot showing time-averaged, relaxed and contraction-phase heart stiffness (two-tailed paired Student’s t-test, \*\*\*\**P* < 0.0001).

The formation of trabeculae is a key step in cardiac development, and disorders of that process lead to cardiomyopathy and embryonic lethality^41,42^. According to image-based simulations, trabeculae act like structural supports that enhance cardiac wall deformability, reduce fluid pressure stresses and homogenise wall stiffness^43^. Their formation is mechanosensitive but emerging models of trabeculation that invoke tissue properties have thus far been based largely on inference from imaging and theory^44–47^. As proof-of-principle of the value of LSE, we mapped trabecular stiffness to generate a morphogenetic hypothesis (Figures 4A-C). An apical to basal stiffness gradient was observed within each trabecula of the E9.25 left ventricle. The stiffness gradient was found to correspond to FN density (Figures 4D and 4E), raising the possibility that myocardial cells move up the stiffness gradient, a process akin to durotaxis, to raise the structure from the cardiac wall (Figure 3F). An increasing number of trabecular cells would increase the production of ECM proteins, potentially feeding back positively upon cell movement into trabeculae until the degree of their stiffness precludes further cell migration similar to what has been observed in limb bud mesoderm^6^. These and other models can be rigorously tested by combining live imaging and genetic approaches with LSE.

**Figure 4.**
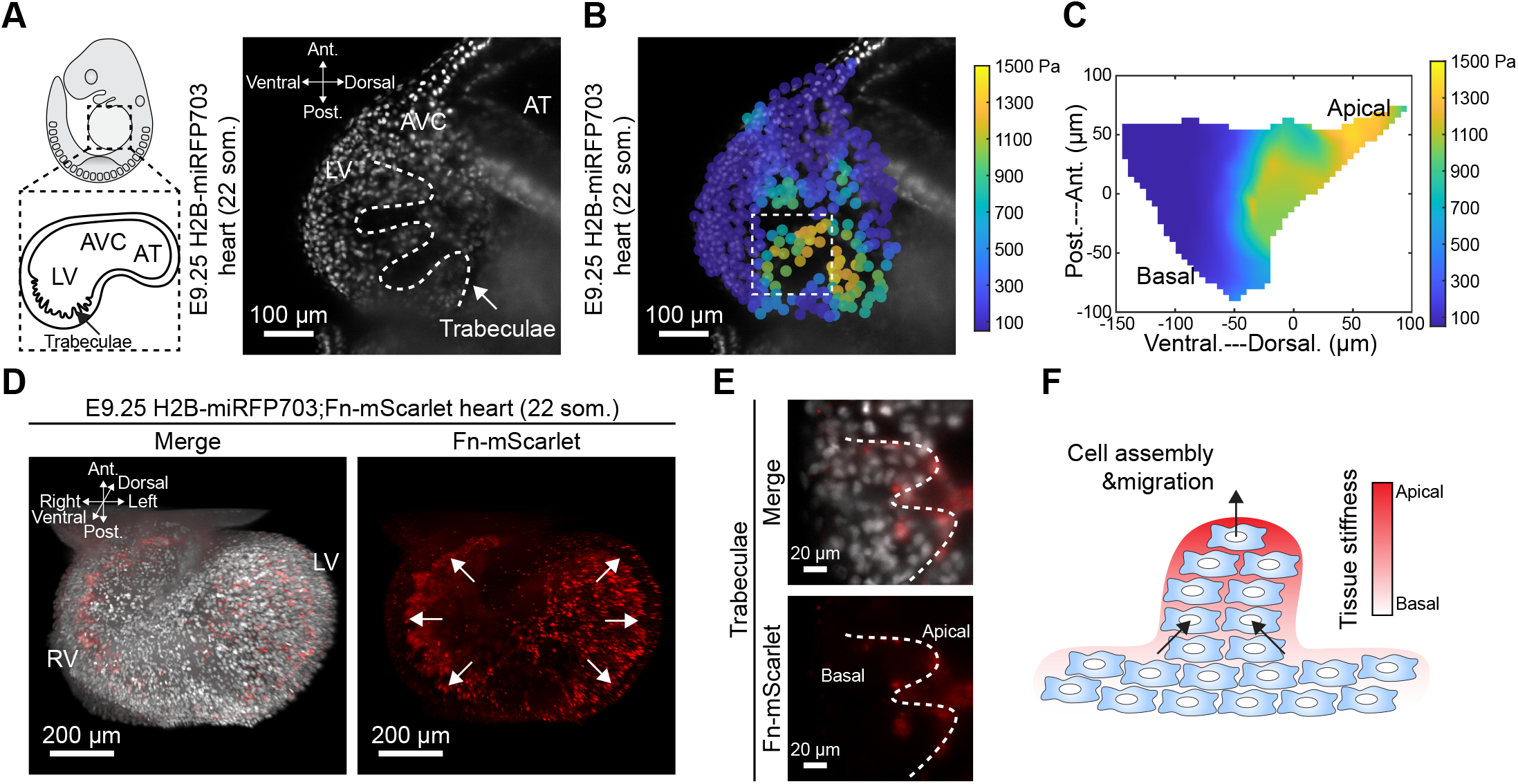
Ventricular traveculation. (**A**) Slice view of an E9.25 H2B-miRFP703 embryonic heart illustrating ventricular trabeculation. (**B**) Trabecular myocardium stiffness map at the focal plane shown in A. (**C**) Rendered zoom-in trabecular myocardium stiffness map (dashed square in B). (**D**) 3D rendering of an E9.25 H2B-miRFP703;Fn-mScarlet embryonic heart. Arrows indicate ventricular trabeculation. (**E**) Zoom-in view of fibronectin expression in trabecular myocardium (dashed line). (**F**) Schematic model representing the potential mechanism that underlies ventricular trabeculation. AT, atria; AVC, atrioventricular canal; LV, left ventricle; RV, right ventricle.

## DISCUSSION

Noninvasively mapping 3D tissue stiffness by LSE offers the potential for a panoramic view of tissue stiffness among various embryos at different stages of development at a spatiotemporal resolution of 15 µm and 0.12 s. With a readily replicable setup, LSE is amenable for integration with a standard light sheet microscope. The noninvasive nature of this method also enables tissue stiffness mapping over time during development. LSE combined with live imaging of molecular reporters permits simultaneous visualisation of biomechanical and biochemical interactions *in vivo*, offering the potential to resolve ‘chicken-and-egg’ questions in morphogenesis. Temporal progression of tissue stiffness also provides valuable inputs for computational modelling and simulation of morphogenetic events. The limitations of LSE stem from the Zeiss Z1 microscope specifications. The Zeiss Z1 can accommodate a sample up to a 1 ml syringe with an inner diameter of 4.76 mm, and its frame rate is limited by a minimum exposure time of 7 ms. While this frame rate is sufficient for mapping embryonic tissue stiffness, a higher frame rate is needed for applications requiring subcellular resolution or imaging of stiff adult tissues which could potentially be achieved by switching to a lattice light-sheet microscope or a customised high-speed light-sheet microscope, respectively^48^.

In cardiac research, tissue stiffness has been invoked but not measured directly *in vivo* due to a lack of quantification tools. LSE’s ability to map stiffness throughout the cardiac cycle and during the morphogenesis of various organ primordia opens avenues for proposing and testing hypotheses concerning development and congenital disease *in vivo* or *in vitro* using organoid models^49^. LSE is compatible with imaging-based mechanical strain mapping^7,50^. Since the product of stiffness and strain yields stress, cardiac tissue stress can be spatiotemporally monitored by computing the product of the stiffness and strain maps. The fast-imaging speed of light sheet microscopy also permits the capture of cardiac conduction velocity (tens of cm/s)^51^. LSE therefore offers the potential for comprehensive cardiac mechanical assessment by simultaneously quantifying cardiac stiffness, stress, and conduction velocity.

An opportune potential application is that LSE makes it possible to register single cell transcriptomic and proteomic datasets spatially with physical properties. Another is that, aside from measurements, actuation of LSE with higher energy offers the potential to manipulate shear stress. Since shear stress can activate signalling cascades that affect morphogenesis^52,53^ and disease progression^54,55^, LSE can be actuated to test mechanotransduction hypotheses.

Our hope is that LSE will render a subset of mechanobiology experiments technically accessible for standard biology labs at low cost.

## MATERIALS AND METHODS

### Animal strains

Analysis was performed using the mouse strain: R26-CAG-H2B-miRFP703 [The Jackson Laboratory: Gt(ROSA)26Sor^em1.1(CAG-RFP*)Jrt^/J], and Fn-mScarlet^6^, outbred to CD1. The zebrafish analysis was performed using Tg(ubi:loxp:mScarlet-nls:loxp:QFGal4)^37^, outbred to TU. All animal experiments were performed in accordance with Canadian Council on Animal Care (CCAC) guidelines, and protocols approved by the Hospital for Sick Children Research Institute Animal Care Committee.

### Light sheet elastography device

The piezoelectric actuator used in this work is P-840.20 (Physik Instrumente). It was attached to the light sheet microscope (Zeiss Lightsheet Z.1) via a mounting stage that was 3D printed using acrylonitrile butadiene styrene (ABS). Three permanent magnets (McMASTER-CARR Lot 5862k178) were used to secure the mounting stage to the Z.1 motorised sample stage. The glass capillary and plunger are standard components for the Z.1. Customised glass capillary holders were 3D printed from resin to connect the glass capillary and actuator. A simple sinusoidal waveform, generated by a function generator (B&K Precision Lot 4053B) and amplified by a custom-made voltage amplifier, was used to drive the piezoelectric actuator. The custom-made voltage amplifier provides a maximum voltage output of 100 V, enabling a travel range of up to 30 μm for the piezoelectric actuator. The design files of the mounting stage, glass capillary holder and voltage amplifier are available for download at https://github.com/MinZhuUOTSickKids/light_sheet_elastography.

### Agarose gel validation

To validate light sheet elastography, we performed stiffness mapping of 0.5 to 3% low melting point agarose gel (ThermoFisher Lot 16520050) in DMEM without phenol red and compared the results to those obtained with AFM indentation. The agarose gel solution contained 5% fluorescent beads of 2 μm diameter (1:200; Sigma Lot L3030 and L4655). The gradient agarose gel was prepared using a diffusion-based method^56^ by mixing agarose of two different percentages containing green and red fluorescent beads, respectively. A 2% agarose gel was aspirated into a glass capillary and solidified at room temperature while a 1% agarose gel solution was kept in a microcentrifuge tube at 95 °C. The glass capillary, containing the two percent agarose gel, was submerged 2 mm into the tube for 20 s, followed by aspiration of the one percent agarose gel. The glass capillary was then maintained in a vertical orientation for agarose gel solidification. Once the agarose solidified, the glass capillary was submerged into the imaging chamber containing DMEM without phenol red, and the agarose plug was partially extruded from the glass capillary. The temperature of the imaging chamber was maintained at 37°C with 5% CO_2_. Upon actuation, the displacements of the fluorescent beads were tracked using Imaris (Bitplane) and fitted in MATLAB (MathWorks) to obtain the phase map. The stiffness map of the agarose gel was resolved from the phase map by computing the phase decay along Y axis.

The stiffness of the same batch agarose gel was also evaluated by AFM (Bruker BioScope Catalyst) mounted on an inverted microscope (Nikon Eclipse-Ti) at 37°C with 5% CO_2_. A spherical probe tip with a diameter of 32 μm was used for all indentations performed. The spherical tip was made by assembling a borosilicate glass microsphere onto a tipless AFM cantilever using epoxy glue. After assembly, the diameter of the probe tip was measured under a scanning electron microscope (Hitachi 4000). The spring constant of the cantilever was calibrated by measuring the power spectral density of the unloaded cantilever thermal noise fluctuation. The linear regime of the approach curve of each indentation was fitted with the Hertz model for a spherical tip to obtain stiffness values^57^.

### Time-lapse tissue stiffness mapping

Dissected embryos were suspended in a solution of DMEM without phenol red containing 12.5% filtered rat serum, 1% low-melt agarose (Invitrogen), and 1% fluorescent beads (1:500, Sigma Lot L3030) that were used for drift-compensation within a glass capillary. Once the agarose solidified, the capillary was submerged into an imaging chamber containing DMEM without phenol red, and the agarose plug was partially extruded from the glass capillary tube until the portion containing the embryo was completely outside of the tube. The temperature of the imaging chamber was maintained at 37°C with 5% CO_2_. Images were acquired for 3 to 4 h at 5 min. intervals.

### Pattern-guided cell motion tracking program

The tracking function in Imaris (Bitplane) is not applicable to the densely populated and fast-moving cells in the LSE embryo dataset as shown in Movie S2. Based on Python, we developed a pattern-guided cell motion tracking program for tracking these cells, utilising a subpixel image registration algorithm^58^. The algorithm compares two images and computes the pixel translation between them by maximising intensity correlation with respect to translation in the two directions.

To evaluate the tracking accuracy, randomly distributed 24 Gaussian blobs of one pixel width each were generated and oscillated with an amplitude similar to LSE actuation (i.e., 15 μm) for 128 time points. The synthetic data were imported into our tracking program and tracking accuracy was quantified by calculating the difference between the real and tracked positions with respect to how far each point moved between each pair of neighbouring time points. The tracking accuracy was determined to be 0.35 pixels permitting the detection of phase shifts as small as 0.0023 rad.

With a user interface, our program allows the selection of a working area to track the average motion. It then tracks selected points within the working area by comparing small areas around each point over time. The points themselves and size of the areas around the points to be tracked are user selectable. The output is the positions of the selected points at each time point. The tracking program and synthetic data generator are available for download at https://github.com/MinZhuUOTSickKids/light_sheet_elastography.

### Spatiotemporal resolution of LSE

The time resolution was determined by fitting varying numbers of frames to the sinusoidal function *A cos* (*ωt* + *φ*). We fitted 3 to 100 frames at an interval of 1 frame. The phase reading was found to stabilize after 7 frames (Figure S5C). To evaluate the spatial resolution of LSE, we quantified the phase root mean squared (RMS) level as a measure of noise, as shown in Figure S5D. The phase RMS is inversely related to time resolution by a power law relation. Considering a signal-to-noise ratio of 3, the minimum phase shift (0.0023 rad at 100 frames as shown in Figure S5D) that can be differentiated by LSE vs. time resolution is

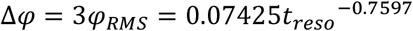

By rewriting the shear modulus - phase shift equation

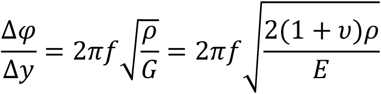

and substituting values in the equation, the spatial resolution of LSE (Δ*y*) in microns is

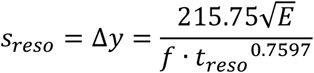

Thus, the spatial resolution at 10 Hz actuation is shown in Figure S5E.

### Apoptosis detection

LysoTracker Red DND-99 (ThermoFisher Lot L7528) was diluted to 2 μM in DMEM containing 50% rat serum. Embryos were placed in media and incubated in a roller culture apparatus for 1 h. The temperature was maintained at 37 °C with 5% CO_2_. Embryos were washed three times with PBS after staining to remove LysoTracker surplus then fixed overnight in 4% paraformaldehyde in PBS followed by three washes for 5 min. in PBS. Images were acquired using the light sheet microscope (Zeiss Lightsheet Z.1) at 20X magnification, and analysis was performed using Imaris.

## Data Availability

The data that support the findings of this study are available at https://figshare.com/articles/dataset/light_sheet_elastography.

## Code Availability

All custom codes used in this paper are available at https://github.com/MinZhuUOTSickKids/light_sheet_elastography.

## ACKNOWLEDGEMENTS

We thank Teng Cui for help with mechanics analysis, and Sepehr Saber for help with DMEM viscosity measurement. This work was funded by the Canada First Research Excellence Fund/Medicine by Design (MbDGQ-2021-04 to SH/YS and MPDF-2020-04 to MZ, and the Canadian Institutes of Health Research (168992) to SH/YS. YS also acknowledges support from the Canada Research Chairs program.

## AUTHOR CONTRIBUTIONS

M.Z. and K.Z. designed and performed the experiments, modelling, and wrote the manuscript; E.T. wrote the tracking program. R.X and B.C. provided zebrafish experimental support. M.Z., Y.S. and S.H. co-conceived the project. Y.S. and S.H. co-funded the project and edited the manuscript.

## SUPPLEMENTARY MOVIE LEGENDS

**Movie S1**

Animation illustrating LSE under ideal pure shear actuation.

**Movie S2**

LSE cell tracking results using Imaris versus our new tracking tool, pattern-guided cell motion tracking (PCMT).

**Movie S3**

LSE actuation on an E9.25 H2B-miRFP703 embryonic heart.

## SUPPLEMENTARY NOTE

Analysis of shear wave propagation in agarose gel

## SUPPLEMENTARY TABLE

Comparison of existing methods and our method in mapping tissue stiffness

**Figure S1.**
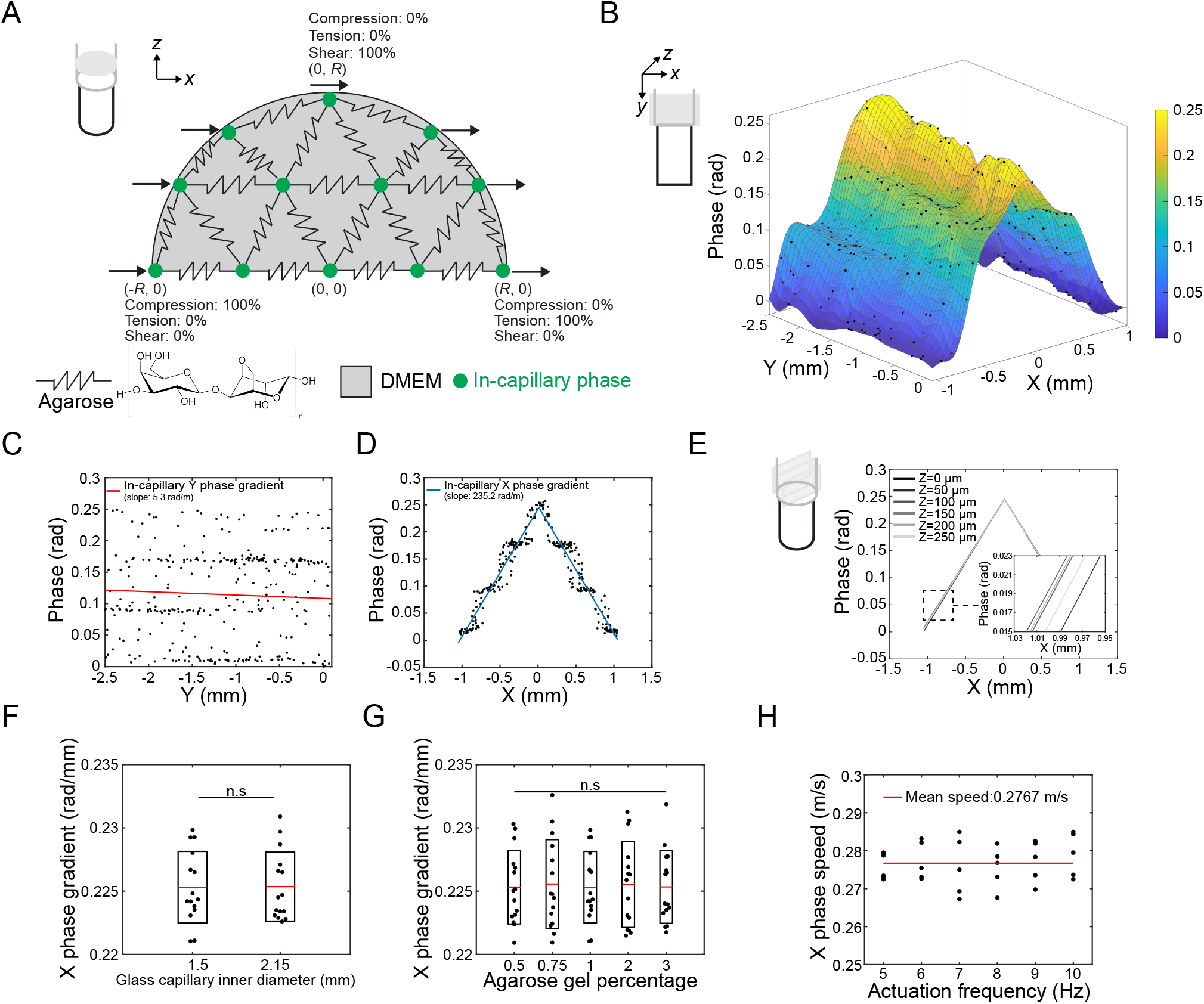
Baseline (X) phase pattern. (A) Schematic depicting the poroelastic agarose gel and boundary conditions at the gel-capillary interface. (B) 3D rendered phase map of 1% agarose gel inside glass capillary under 10 Hz actuation. (C and D) Glass capillary phase along Y axis (C) and X axis (D). (E-G) X phase gradient is independent of the Z location (E), glass capillary diameter (F), and agarose gel percentage (G). (n=3 for each diameter and gel percentage, two-tailed unpaired Student’s t-test). n.s, not significant. (H) Constant X phase speed under different actuation frequencies suggesting a non-dispersive X wave.

**Figure S2.**
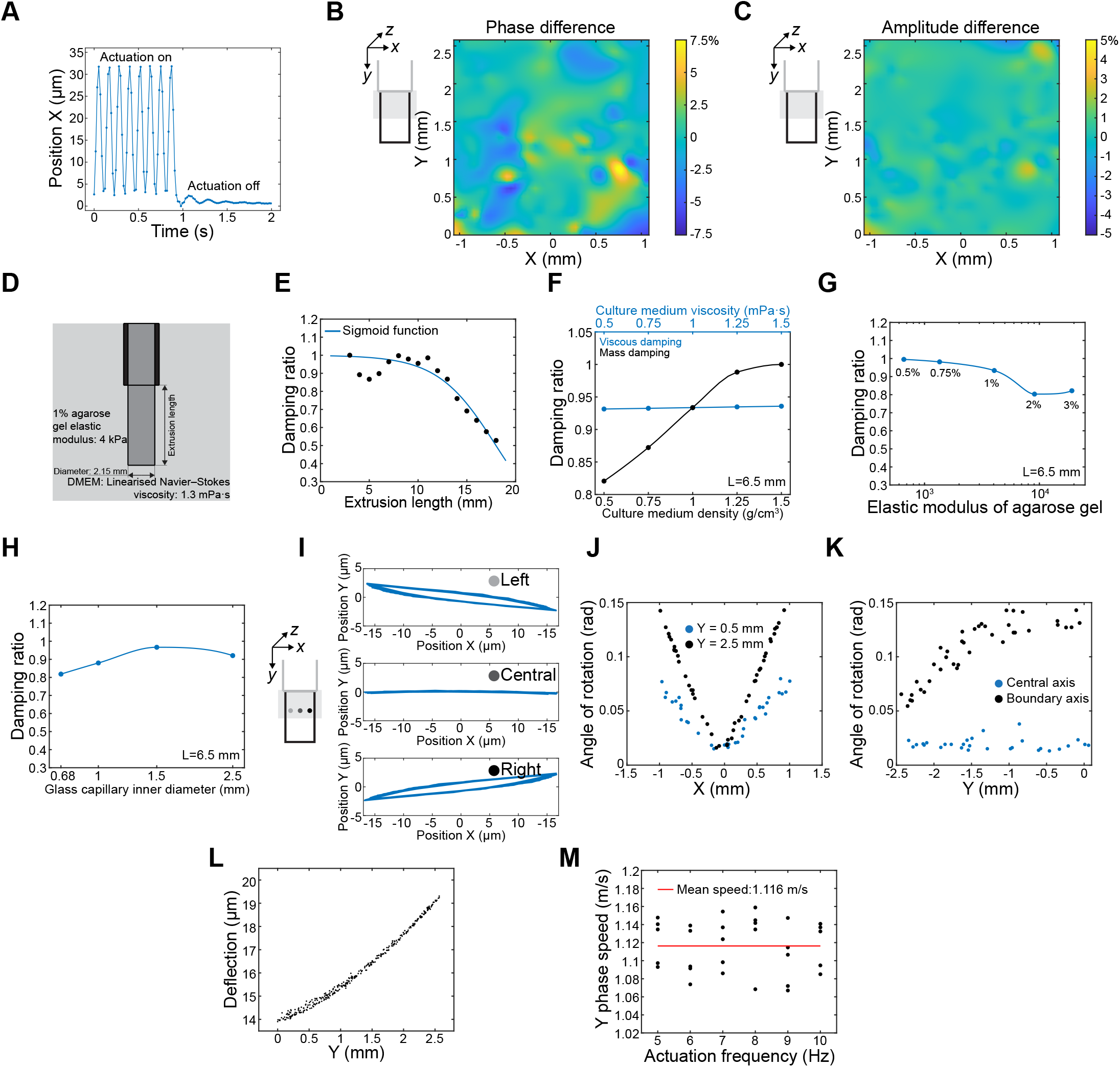
Y phase pattern. (**A**) Representative position tracking of a fluorescent bead in 1% agarose gel upon 10 Hz actuation. (**B** and **C**) 3D rendered phase and amplitude difference submerged in DMEM vs water at 37°C. (**D**) Fluid-structure interaction simulation model setup. (**E**) Simulation-predicted damping ratio as a function of the gel extrusion length. (**F**) Simulation-predicted damping ratio at varying viscosity and culture medium density with an extrusion length of 6.5 mm suggesting predominantly mass damping. (**G** and **H**) Simulation -predicted damping ratio at various agarose gel elastic moduli (G) and diameters (H) with an extrusion length of 6.5 mm. (**I** and **J**) Angle of rotation along central-boundary axis. (**K**) Angle of rotation along Y axis. (**L**) Increased deflection along Y axis. (**M**) Constant Y phase speed under different actuation frequencies suggesting non-dispersive shear wave propagation along the Y axis.

**Figure S3.**
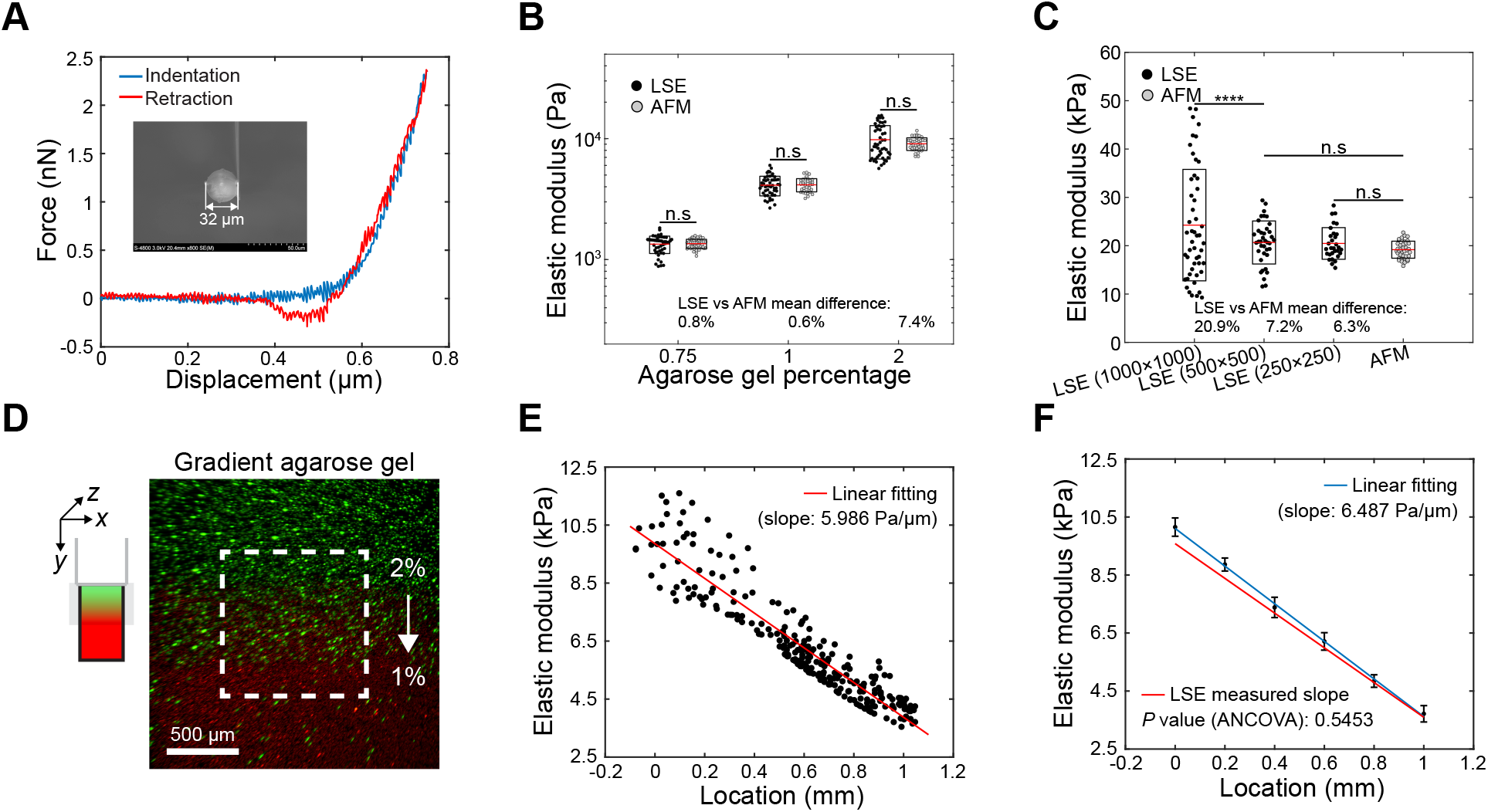
AFM validation. (**A**) Representative AFM indentation-retraction curve without hysterisis suggesting elastic behaviour of the agarose gel (inset: scanning electron micrograph of an AFM cantilever with a spherical tip of 32 μm diameter). (**B**) Comparison of agarose gels (0.5% to 2%) with distinct elastic moduli measured by LSE and AFM (n=3 for each gel percentage, two-tailed unpaired Student’s t-test). (**C**) 3% agarose gel measured by LSE and AFM illustrating lowering CMOS size from 1000×1000 (default) to 500×500 increase LSE measurable stiffness range at the cost of field-of-view (n=3, two-tailed unpaired Student’s t-test, \*\*\*\**P* < 0.0001). (**D**) Representative image illustrating 2% to 1% gradient agarose gel that contains green and red fluorescent beads, respectively. (**E** and **F**) 2% to 1% agarose gel stiffness measured by LSE (E) and AFM (F) showing no statistical difference in the stiffness gradient. n.s, not significant. Error bars indicate s.d..

**Figure S4.**
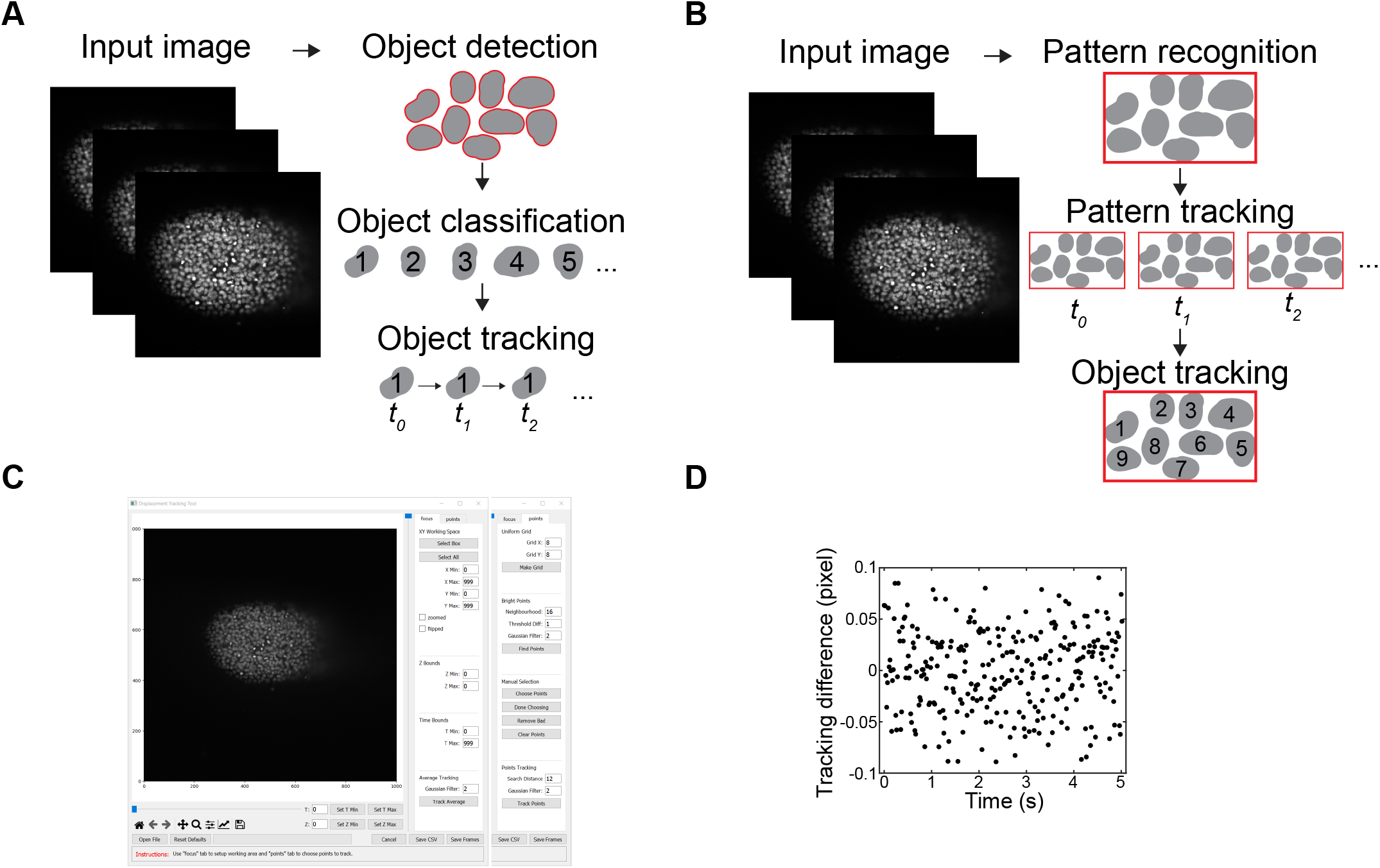
Pattern-guided cell motion tracking program. (**A**) Schematic describing the autoregressive motion algorithm used in Imaris. (**B**) Schematic describing the tracking algorithm used in pattern-guided cell motion tracking program. (**C**) GUI of the pattern-guided cell motion tracking program. Neighbourhood represents the density of the cells. (**D**) Imaris and PCMT sub-pixel level tracking difference of a fluorescent bead embedded in 1% agarose gel under 10 Hz actuation.

**Figure S5.**
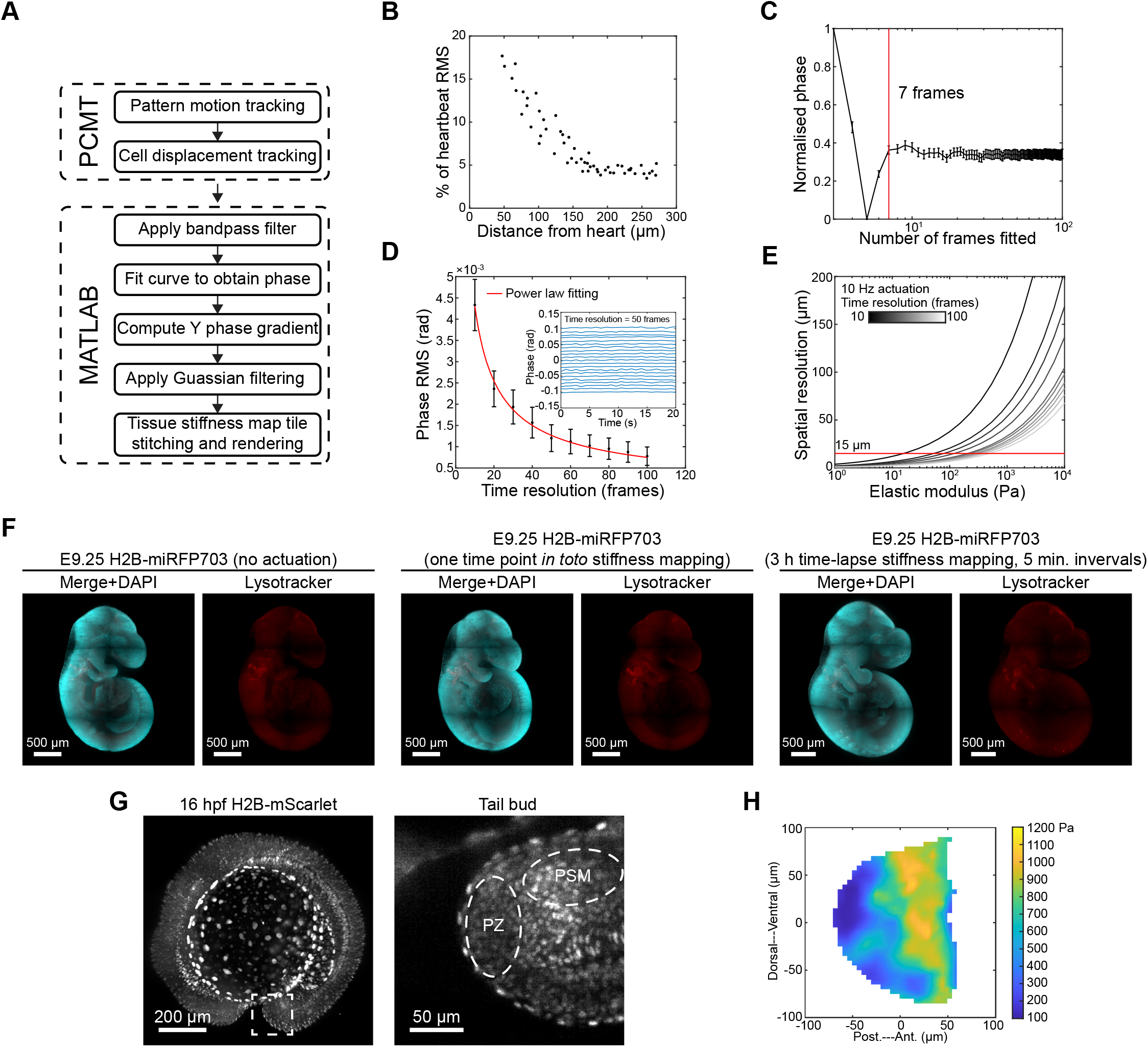
Non-invasive tissue stiffness mapping. (**A**) Flowchart illustrating LSE analysis pipeline. (**B**) Heart beat effect decays over distance. (**C**) LSE time resolution. (**D**) Phase root mean square (RMS) as a function of time resolution (inset: examples of phase fluctuation over time at 50 frames time resolution). (**E**) LSE spatial resolution as a function of time resolution, actuation frequency and sample stiffness. (**F**) 3D rendering of E9.25 H2B-miRFP embryos stained with DAPI (cyan) lysotracker (red) under no actuation (left panel), one time point *in toto* stiffness mapping (middle panel), and 3 h time-lapse stiffness mapping at 5 min. intervals (right panel) showing no significant increase in cell apoptosis (n=3 for each condition). (**G**) 3D rendering of a 16 hpf H2B-mScarlet zebrafish embryo. Right panel showing a zoomed-in view of the tail bud. PZ, progenitor zone; PSM, presomitic mesoderm. (**H**) Tail bud (right panel in G) stiffness map showing an anterior to posterior stiffness gradient. Error bars indicate s.d..

## SUPPLEMENTARY NOTE

Analysis of shear wave propagation in agarose gel

A prerequisite for using shear wave propagation to quantify stiffness was to establish a phase-shear wave propagation speed relation. The complexity of the multi-physical interactions within the system affirmed the impracticality of an analytical approach. Therefore, we strategised our analysis by identifying and examining the contributing factors of the local phase to arrive at an empirical solution for local phase. For this purpose, we discuss: 1) general wave equation, 2) material behaviour under actuation, 3) baseline phase pattern, 4) propagation of longitudinal waves, 5) propagation of shear waves, 6) culture medium damping, and 7) geometry considerations. A summary of these contributing factors was illustrated in Figure 1D.

### 1. General wave equation

For a one-dimensional system under the influence of propagating waves along X induced by an external force, the general form of the one-dimensional wave equation is[1]

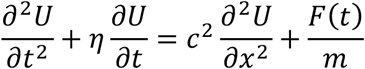

where *c* is the wave propagation speed, and damping coefficient is denoted as *η*. For waves propagating in the absence of damping, the above equation simplifies to

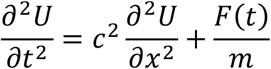

The general solution is,

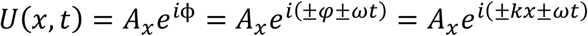

where ϕ is phase angle, *φ* is local phase due to wave propagation, and *k* is wavenumber.

### 2. Material behaviour under actuation

Characteristics of the excitation signal underpin the baseline behaviour of the system. Since the piezoelectric actuator was driven by a sinusoidal signal, the displacement function, *U*_0_(*t*), takes on the form,

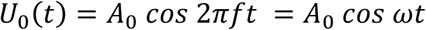

where *A*_0_ is the amplitude of the signal, *f* is the temporal actuation frequency, and *ω* is the angular actuation frequency. To achieve optimal tracking and curve fitting results, it is important to maintain an adequate amplitude while avoiding excessive rotational bending and deflection (refer to the geometry considerations section). We conducted an experimental parametric sweep of the amplitude from 5-15 μm with a step size of 1 μm. An amplitude above 10 μm provided a good fit, and in the 10-15 μm range, the R-squared value was above 0.98. The actuation frequency was selected based on the microscope settings and viscoelastic behavior of agarose gel. In a typical 1000 × 1000 pixels setting, the Zeiss Lightsheet Z.1 gives a frame rate of 57.2 Hz limiting the maximum actuation frequency to 28.6 Hz according to the Nyquist-Shannon sampling theorem. To mitigate the viscoelastic behaviour of the agarose gel for a simple solution of phase, an actuation frequency sweep from 5 to 25 Hz with an amplitude of 15 μm was conducted. We found that actuation frequencies under 10 Hz yield good tracking results, with the phase speed being independent of frequency in both the X and Y axes, suggesting elastic behaviour (Figures S1H and S2M). From the AFM indentation experiment performed under same strain rate estimated by simulation, no indent-retract hysteresis was observed, which further validated the predominant elastic behaviour of agarose gel as shown in Figure S3A.

When a structure undergoes forced oscillation, the actuation induces a collection of local interactions among the particles within the structure. The local displacement pattern is determined by direct actuation and other system-specific factors. At steady state, the generalised displacement of each particle located at (*x, y*) takes the following form[1],

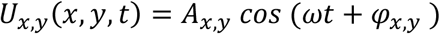

where *φ*_*x,y*_ is the local phase. The phase of the extruded agarose gel consists of two parts: the shear wave propagation phase along the Y-axis and the phase inherited from the agarose gel inside the glass capillary, henceforth referred to as the baseline phase.

### 3. Baseline phase pattern

To establish a baseline phase pattern, we conducted sinusoidal actuation and tracked the displacement of fluorescent beads embedded in the agarose gel. The field-of-view used was 2.6 mm × 2.6 mm covering the entire agarose gel within the glass capillary. The displacement of an individual bead was tracked by weighted pixel intensity and fitted with a sinusoidal function to obtain the local phase *φ*_*x,y*_. For each actuation, over 300 fluorescent beads were tracked and fitted with R-squared values above 0.98 to generate the baseline phase pattern as shown in Figure S1B. The baseline phase pattern displayed a symmetrical decay along X folded at the center of the capillary without changes in phase along Y (Figures S1C and S1D). Multiple ruffle folds were located along X at a phase interval of ∼0.075 rad. The X phase gradient was independent of Z location, glass capillary diameter, and stiffness (i.e., agarose gel concentration) as shown in Figures S1E-S1G. The X phase speed (i.e., wave propagation speed) was found to be constant at different actuation frequencies suggesting the wave being non-dispersive. The X phase pattern results from a complex interplay of boundary conditions, including adhesion and decohesion between the gel and glass capillary, as well as wave propagation, reflection, and interference.

### 4. Longitudinal wave propagation

Longitudinal wave propagation contributes to displacement along the X axis inside the glass capillary. Based on the general wave equation, the displacement due to longitudinal wave propagation without viscous damping effect can be expressed as,

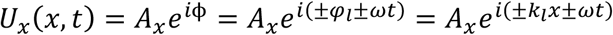

where ϕ is the phase angle, *φ*_*l*_ is the local phase due to longitudinal wave propagation, and *k*_*l*_ is the longitudinal wavenumber. If only the real-time forward-moving longitudinal wave is considered, the horizontal displacement pattern of a particle at X can be expressed as,

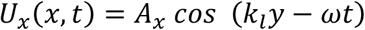

The local phase due to longitudinal wave propagation *φ*_*l*_ is a function of the horizontal location of the particles, i.e.,

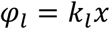

The longitudinal wavenumber *k*_*l*_ can be used to calculate the propagation speed of the longitudinal wave *c*_*l*_, which is determined by the longitudinal modulus *M* and density *ρ* of the material by,

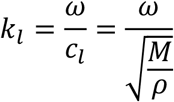

Longitudinal wave propagates at a speed around 1,500 m/s in agarose gels of different concentrations[2] which corresponds to a phase gradient along X axis, Δ*φ*_*l*_/Δ*x* = 0.0016 rad/mm (less than 1% of the observed X phase gradient in the capillary) under 10 Hz actuation. The phase delay caused by longitudinal wave propagation can therefore be neglected from our phase analysis. The longitudinal wave propagation speed in agarose gel was discovered to be dominated by the fluid component which constitutes over 90% of its mass and largely independent of gel stiffness (i.e., concentrations)[2]. Considering the insufficient imaging frame rate offered by Zeiss Z.1 (i.e., up to 100 fps) to capture the rapid propagation, decoupling the longitudinal modulus of the gel from its propagation speed is infeasible.

### 5. Shear wave propagation

In the extruded gel, the horizontal motion of particles located along Y at a fixed X can be perceived as a 1D system under the influence of Y-propagating shear waves. Based on the general wave equation, the displacement due to shear wave propagation without damping effect can be expressed as,

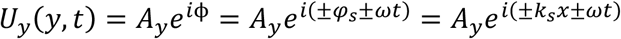

where ϕ is the phase angle, *φ*_*s*_ is the local phase due to shear wave propagation, and *k*_*s*_ is the shear wavenumber. If only the real-time forward-moving shear wave is considered, the horizontal displacement pattern of a particle at Y can be expressed as,

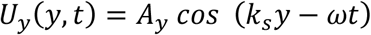

The local phase due to shear wave propagation *φ*_*s*_ is a function of the vertical location of the particles, i.e.,

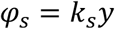

The shear wavenumber (*k*_*s*_) can be used to quantify the shear modulus (*G*) of the material via,

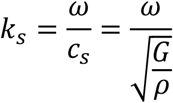

Unlike longitudinal waves, shear waves propagate at a much lower speed in agarose gels (a few m/s)[3], making it ideal for decouple the shear modulus of the gel.

### 6. Culture medium damping

The medium in which the gel was submerged, DMEM, served multiple purposes. First, DMEM acted as a culture medium that supported the viability and growth of a living biological sample. Second, DMEM has a similar density to agarose gel, thus counterbalancing the effect of gravity loading on the actuated gel through buoyancy. DMEM also served as an external damping source to minimize wave reflections and gel rotational bending and deflection induced by the actuation. DMEM damping can be separated into two parts, the added mass of the fluid that is displaced by the agarose gel (i.e., mass damping) and the viscous fluid shear at the surface of the gel (viscous damping).

To investigate the effect of DMEM damping on local phase, we considered DMEM as an infinite medium since the size of the chamber is orders of magnitudes larger than the actuation amplitude. Based on the general wave equation, 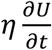, was introduced. The solution for a damped wave acquired a complex form (*Ũ*) that can be expressed as follows[1],

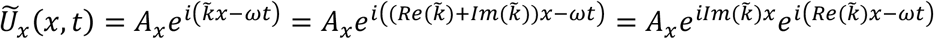

The real term of the solution became,

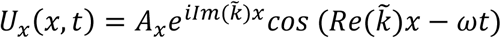

The complex wavenumber 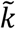 consisted of a real part, 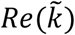, and an imaginary part, 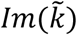, whose expressions take the following forms,

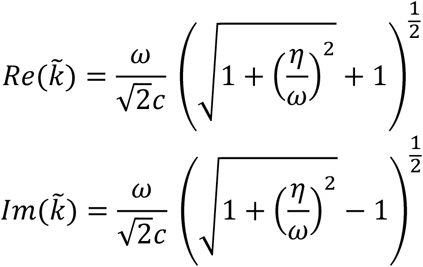

As a result, adding a damping component affects both local amplitude and phase. The magnitude to which their values are altered depends on the damping coefficient *η*. The damping ratio *ζ* relates *η* and undamped natural frequency of the system *ω*_*n*_ by,

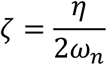

The damping ratio *ζ* can vary from undamped (*ζ* =0), underdamped (*ζ* <1), critically damped (*ζ* =1) to overdamped (*ζ* >1). Experimental observations indicated that our system was closely approaching critical damping by DMEM as illustrated in Figure S2A. To investigate the impact of system parameters (e.g., gel extrusion length) on the damping behavior and the source of DMEM-induced damping (i.e., mass vs. viscous damping), we conducted COMSOL fluid-structure interaction (FSI) simulations. The inputs of the simulation are illustrated in Figure S2D. DMEM was modelled as linearized Navier-Stokes with a dynamic viscosity of 1.3 mPa·s measured using a rheometer (TA Instruments, AR 2000) at 37 °C. The input elastic modulus for 1% agarose was 4.05 kPa determined by AFM indentation at 37 °C. We simulated 1% agarose gel with an extrusion length ranging from 3-18 mm, given the fact that 3 mm being the minimum length required to properly fit an E9.25 mouse embryo whereas 18 mm being the maximum length without interfering with the imaging chamber. For extrusion lengths less than 12 mm, the damping ratio clustered around 1, suggesting a critical damping state. As the extrusion exceeded 12 mm, the damping ratio continued to decrease, moving the system into an underdamped state. By sweeping the culture medium viscosity (from 0.5 to 1.5 mPa·s) and density (0.5 to 1.5 g/cm^3^) in the FSI simulation, we found that damping ratio is predominantly determined by density (Figure S2F). Since various culture media have densities similar to that of water, replacing the culture medium with different viscosity in the LSE experiment does not affect the critical damping conclusion. This was experimentally validated by substituting DMEM with water (viscosity: 0.69 mPa·s at 37 °C), as depicted in Figures S2B and S2C. By sweeping through agarose gel elastic moduli (0.5% to 3%, as determined by AFM) and four stock Zeiss Z.1 glass capillaries of different inner diameters, critical damping effect was verified for different gel percentage and diameter (Figures S2G and S2H).

In summary, for agarose gel with an extrusion length spanning from 3 to 12 mm, the system can be regarded as being critically damped. The critical damping effect, dominated by the density of culture medium, filters out undesired wave reflection and mitigates its contribution to the local phase under forced oscillation[1]. As a result, the damping effect can be disregarded for phase analysis. It is noted that, as the dimensions of the agarose gel (diameter and extrusion length) decrease to the micrometer level (e.g., AFM cantilever), the influence of viscous damping becomes significant and must be taken into consideration[4].

### 7. Geometric considerations

For the analysis of shear wave propagation, agarose gel was simplified as a one-dimensional system. The loaded three-dimensional structure of agarose gel imposes rotational bending and deflection governed by beam vibration theories. Understanding the particle’s response to local rotational bending and deflection is necessary for the phase evaluation as well as the design of the post-processing algorithm. Under a dynamic load, *F*(*t*), the characteristics of a homogenous, isotropic, linearly elastic thin beam with a uniform transverse cross-section XZ under a small deformation can be described by Timoshenko-Ehrenfest beam theory (TEBT). The horizontal displacement pattern *U* is[5],

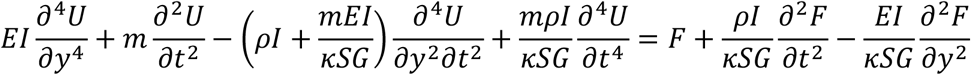

with *I* representing the second moment of area of the structure, *m* representing the mass of the structure, *S* representing the transverse cross-sectional area, and *κ* representing the Timoshenko shear coefficient that depends on the cross-sectional geometry. For a solid beam with a circular transverse cross-section, *κ* is 0.9[6]. The dynamic load introduced a rigid movement, with the time-dependent terms representing a combined effect of inertia and shear forces concerning local rotation as a function of Y, and the fourth-order partial derivative with respect to Y characterizing local deflection. Considering the minimal damping effect from DMEM and no sliding motion between the gel and glass capillary, our system can be modelled as a free-end at *y* = *L* and a fix-end at *y* = 0.

A low actuation frequency results in an actuation wavelength longer than the length of the extruded gel. For 1% agarose gel under 10 Hz actuation, the actuation wavelength was 1.16 m while the extrusion length of the gel was less than 12 mm. The system resembles a beam undergoing first-mode antisymmetric flexural vibration[7]. The portion to one side of the central axis (*x* = 0) would be under tension while the other side be under compression at an arbitrary time point and vice versa. The macroscopic distortion of the geometry can be decomposed into local rotation at the single-particle level. To study the characteristics of local rotation, the Timoshenko beam equation is rewritten in its equivalent coupling form by directly integrating the angle of rotation,

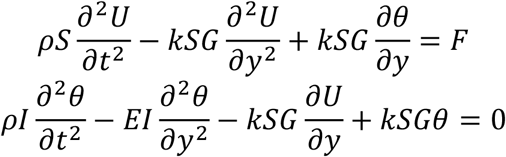

The rotational effect can be understood through the angle of rotation, *θ*, as a function of the particle location along *y*. For a particle located at *y*, the 2D displacement field vector *D* is composed of the horizontal *U* and vertical displacement *V* vectors by[8],

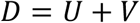

The angle of rotation is defined as,

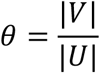

The local rotation induces an elliptically polarized motion instead of a pure horizontal motion (X direction) as shown in Figure S2I. We next examined the local rotation along the X and Y axes. Along the X axis, minimum rotation was observed near the central axis. A linear relation was found between the horizontal distance from the central axis and angel of rotation (Figure S2J). Along the Y axis, the maximum angle of rotation at boundary increased with Y location but plateaued at 0.15 rad as shown in Figure S2K. Therefore, the displacement of particles was approximated as *U*.

For local deflection, the inertial and shearing effects can be excluded, resulting in a simplified form of TEBT known as the Euler-Bernoulli beam theory (EBBT) that can be expressed as[9],

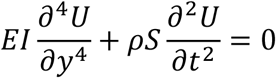

Premised on the given boundary conditions, the fix-end adopts the displacement function, *U*_0_(*t*), as prescribed by the piezoelectric actuator with no Y dependence. The free-end is assumed to be without rotational or shearing forces. The boundary conditions can be expressed as,

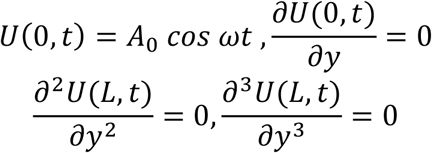

The solution is,

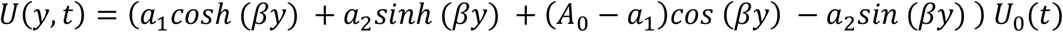

with the coefficients as,

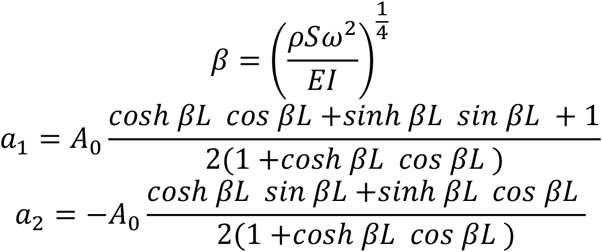

The solution of EBBT suggests an increase in the displacement amplitude towards +Y which was confirmed by experimental results (Figure S2L). The increase in displacement amplitude does not affect the local phase since *U* and *U*_0_ remain in phase when located away from the node. The increased local bead movement speed induced by the increased displacement amplitude undermines tracking accuracy. As such, to achieve optimal tracking results, we limited the maximum actuation frequency to 10 Hz.

Conclusion

During light sheet live imaging, considering that longer gel extrusion length can cause tissue drift over time, we selected extrusion length from 6 to 7 mm for our tissue stiffness mapping experiments with a maximum actuation frequency of 10 Hz for optimal tracking results. As analysed in previous sections, the phase relationship can be empirically solved as,

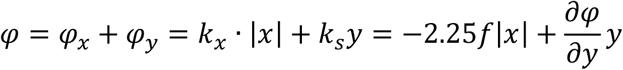

where *k*_*x*_ and is *k*_*s*_ are X and Y wavenumbers, respectively.

## SUPPLEMENTARY TABLE

Comparison of existing methods and our method in mapping tissue stiffness

| Method | AFM | Magnetic and optical tweezers | Brillouin microscopy | C-OCE | Clinical elastography | Light sheet elastography |
| --- | --- | --- | --- | --- | --- | --- |
| Measurement | 2D | 3D single point or multiple points | 3D mapping | 3D mapping | 2D and 3D mapping | 3D mapping |
| Invasiveness | Noninvasive for surface measurement | Invasive | Phototoxicity | Noninvasive | Noninvasive | Noninvasive |
| Assumption | Hertzian contact | Mechanics model (e.g., standard linear solid) | Tissue refractive index and density | Tissue refractive index and uniform stress distribution | Tissue density | Tissue density |
| Spatial resolution | Subcellular | Subcellular (cellular for multi-beads actuation) | Subcellular | Subcellular | mm to cm | Cellular |
| Time resolution | ms to s per point | ms to s per point | 20-500 ms per pixel | 3-6 min per volume | ms to min | 0.12 s per 1000 × 1000 pixel image |
| Validation | Often taken as gold standard | Cross validated by AFM | Positive and no correlation both reported when benchmarked against AFM | Cross validated by standard compression test | Cross validated by tensile test | Cross validated by AFM |
| Major limitation | Invasive when measuring underlying tissues | Invasive; spatial resolution limited by number of injected beads | Slow acquisition for multi-slice 3D tissue stiffness mapping | Error caused by stress variation in geometrically complex tissues and limited axial resolution of strain measurement (~ 300 $\mu\text{m}$) | Limited spatial resolution | Sample size limited by Zeiss Z1 (1 ml syringe; inner diameter: 4.76 mm) |
| Example | [10], [11] | [12], [13] | [14], [15] | [16], [17] | [18], [19] | present manuscript |

## REFERENCE

1. Mongera, A. et al. A fluid-to-solid jamming transition underlies vertebrate body axis elongation. Nature Preprint at 10.1038/s41586-018-0479-2 (2018).

2. Barriga, E. H., Franze, K., Charras, G. & Mayor, R. Tissue stiffening coordinates morphogenesis by triggering collective cell migration in vivo. Nature 554, 523–527 (2018).

3. Zhu, M. et al. Spatial mapping of tissue properties in vivo reveals a 3D stiffness gradient in the mouse limb bud. Proc Natl Acad Sci U S A (2020) doi:10.1073/pnas.1912656117.

4. Hannezo, E. & Heisenberg, C. P. Mechanochemical Feedback Loops in Development and Disease. Cell 178, 12–25 (2019).

5. Parada, C. et al. Mechanical feedback defines organizing centers to drive digit emergence. Dev Cell 57, 854-866.e6 (2022).

6. Zhu, M. et al. A fibronectin gradient remodels mixed-phase mesoderm. Sci Adv 10, (2024).

7. Tao, H. et al. Oscillatory cortical forces promote three dimensional cell intercalations that shape the murine mandibular arch. Nat Commun (2019) doi:10.1038/s41467-019-09540-z.

8. Serwane, F. et al. In vivo quantification of spatially varying mechanical properties in developing tissues. Nat Methods 14, 181–186 (2017).

9. D’Angelo, A., Dierkes, K., Carolis, C., Salbreux, G. & Solon, J. In Vivo Force Application Reveals a Fast Tissue Softening and External Friction Increase during Early Embryogenesis. Current Biology (2019) doi:10.1016/j.cub.2019.04.010.

10. Zhu, M., Zhang, K., Tao, H., Hopyan, S. & Sun, Y. Magnetic Micromanipulation for In Vivo Measurement of Stiffness Heterogeneity and Anisotropy in the Mouse Mandibular Arch. Research (2020) doi:10.34133/2020/7914074.

11. Bambardekar, K., Clément, R., Blanc, O., Chardès, C. & Lenne, P.-F. Direct laser manipulation reveals the mechanics of cell contacts in vivo. Proceedings of the National Academy of Sciences (2015) doi:10.1073/pnas.1418732112.

12. Zhang, K., Zhu, M., Thomas, E., Hopyan, S. & Sun, Y. Existing and Potential Applications of Elastography for Measuring the Viscoelasticity of Biological Tissues In Vivo. Front Phys 9, 670571 (2021).

13. Prevedel, R., Diz-Muñoz, A., Ruocco, G. & Antonacci, G. Brillouin microscopy: an emerging tool for mechanobiology. Nature Methods 2019 16:10 16, 969–977 (2019).

14. Kabakova, I. et al. Brillouin microscopy. Nature Reviews Methods Primers 4, (2024).

15. Scarcelli, G. et al. Noncontact three-dimensional mapping of intracellular hydromechanical properties by Brillouin microscopy. Nat Methods (2015) doi:10.1038/nmeth.3616.

16. Schlüßler, R. et al. Mechanical Mapping of Spinal Cord Growth and Repair in Living Zebrafish Larvae by Brillouin Imaging. Biophys J 115, 911–923 (2018).

17. Martinez-Vidal, L. et al. Progressive alteration of murine bladder elasticity in actinic cystitis detected by Brillouin microscopy. Scientific Reports 2024 14:1 14, 1–16 (2024).

18. Möllmert, S. et al. Beyond comparison: Brillouin microscopy and AFM-based indentation reveal divergent insights into the mechanical profile of the murine retina. bioRxiv 2024.01.24.577013 (2024) doi:10.1101/2024.01.24.577013.

19. Schlüßler, R. et al. Correlative all-optical quantification of mass density and mechanics of sub-cellular compartments with fluorescence specificity. Elife 11, (2022).

20. Wu, P. J. et al. Water content, not stiffness, dominates Brillouin spectroscopy measurements in hydrated materials. Nature Methods 2018 15:8 15, 561–562 (2018).

21. Koski, R. 1 et al. Reply to ‘Water content, not stiffness, dominates Brillouin spectroscopy measurements in hydrated materials’. Nature Methods 2018 15:8 15, 562–563 (2018).

22. Zaitsev, V. Y. et al. Strain and elasticity imaging in compression optical coherence elastography: The two-decade perspective and recent advances. J Biophotonics 14, e202000257 (2021).

23. Ophir, J., Céspedes, I., Ponnekanti, H., Yazdi, Y. & li, X. Elastography: A Quantitative Method for Imaging the Elasticity of Biological Tissues. 10.1177/016173469101300201 13, 111–134 (1991).

24. Schmitt, J. M. OCT elastography: imaging microscopic deformation and strain of tissue. Optics Express, Vol. 3, Issue 6, pp. 199–211 3, 199–211 (1998).

25. Kennedy, K. M. et al. Quantitative micro-elastography: imaging of tissue elasticity using compression optical coherence elastography. Scientific Reports 2015 5:1 5, 1–12 (2015).

26. Kennedy, K. M., Ford, C., Kennedy, B. F., Bush, M. B. & Sampson, D. D. Analysis of mechanical contrast in optical coherence elastography. J Biomed Opt 18, (2013).

27. Yoshida, S. Waves: Fundamentals and dynamics. Waves: Fundamentals and dynamics (2017) doi:10.1088/978-1-6817-4573-2.

28. Tang, A., Cloutier, G., Szeverenyi, N. M. & Sirlin, C. B. Ultrasound elastography and MR elastography for assessing liver fibrosis: Part 1, principles and techniques. American Journal of Roentgenology 205, 22–32 (2015).

29. Loomba, R. et al. Novel 3D Magnetic Resonance Elastography for the Noninvasive Diagnosis of Advanced Fibrosis in NAFLD: A Prospective Study. American Journal of Gastroenterology 111, 986–994 (2016).

30. Grasland-Mongrain, P. et al. Ultrafast imaging of cell elasticity with optical microelastography. Proc Natl Acad Sci U S A 115, (2018).

31. Gu, B., Posfai, E. & Rossant, J. Efficient generation of targeted large insertions by microinjection into two-cell-stage mouse embryos. Nat Biotechnol (2018) doi:10.1038/nbt.4166.

32. Zörner, S., Kaltenbacher, M., Lerch, R., Sutor, A. & Döllinger, M. Measurement of the elasticity modulus of soft tissues. J Biomech 43, 1540–1545 (2010).

33. Walker, J. M. et al. Nondestructive evaluation of hydrogel mechanical properties using ultrasound. Ann Biomed Eng 39, 2521–2530 (2011).

34. McDole, K. et al. In Toto Imaging and Reconstruction of Post-Implantation Mouse Development at the Single-Cell Level. Cell (2018) doi:10.1016/j.cell.2018.09.031.

35. Thurman, S. T., Guizar-Sicairos, M. & Fienup, J. R. Efficient subpixel image registration algorithms. Optics Letters, Vol. 33, Issue 2, pp. 156–158 33, 156–158 (2008).

36. Taljanovic, M. S. et al. Shear-Wave Elastography: Basic Physics and Musculoskeletal Applications. 10.1148/rg.2017160116 37, 855–870 (2017).

37. Burgess, J. et al. An optimized QF-binary expression system for use in zebrafish. Dev Biol 465, 144–156 (2020).

38. Lau, K. et al. Anisotropic stress orients remodelling of mammalian limb bud ectoderm. Nat Cell Biol 17, 569–579 (2015).

39. Bi, D., Lopez, J. H., Schwarz, J. M. & Manning, M. L. A density-independent rigidity transition in biological tissues. Nature Physics 2015 11:12 11, 1074–1079 (2015).

40. Martin, J. A., Schmitz, D. G., Ehlers, A. C., Allen, M. S. & Thelen, D. G. Calibration of the shear wave speed-stress relationship in ex vivo tendons. J Biomech 90, 9–15 (2019).

41. Captur, G. et al. Morphogenesis of myocardial trabeculae in the mouse embryo. J Anat 229, 314–325 (2016).

42. del Monte-Nieto, G. et al. Control of cardiac jelly dynamics by NOTCH1 and NRG1 defines the building plan for trabeculation. Nature 2018 557:7705 557, 439–445 (2018).

43. Cairelli, A. G. et al. Role of tissue biomechanics in the formation and function of myocardial trabeculae in zebrafish embryos. J Physiol 602, 597–617 (2024).

44. Cho, J. M. et al. Quantitative 4D imaging of biomechanical regulation of ventricular growth and maturation. Curr Opin Biomed Eng 26, 100438 (2023).

45. Taber, L. A. & Zahalak, G. I. Theoretical Model for Myocardial Trabeculation. (2001) doi:10.1002/1097-0177.

46. de Boer, B. A., Le Garrec, J. F., Christoffels, V. M., Meilhac, S. M. & Ruijter, J. M. Integrating multi-scale knowledge on cardiac development into a computational model of ventricular trabeculation. Wiley Interdiscip Rev Syst Biol Med 6, 389–397 (2014).

47. Cairelli, A. G., Chow, R. W. Y., Vermot, J. & Yap, C. H. Fluid mechanics of the zebrafish embryonic heart trabeculation. PLoS Comput Biol 18, e1010142 (2022).

48. Patel, K. B. et al. High-speed light-sheet microscopy for the in-situ acquisition of volumetric histological images of living tissue. Nature Biomedical Engineering 2022 6:5 6, 569–583 (2022).

49. Lewis-Israeli, Y. R. et al. Self-assembling human heart organoids for the modeling of cardiac development and congenital heart disease. Nature Communications 2021 12:1 12, 1–16 (2021).

50. Gjorevski, N. & Nelson, C. M. Mapping of Mechanical Strains and Stresses around Quiescent Engineered Three-Dimensional Epithelial Tissues. Biophys J 103, 152–162 (2012).

51. Weber, M. et al. Cell-accurate optical mapping across the entire developing heart. Elife 6, (2017).

52. Steed, E. et al. klf2a couples mechanotransduction and zebrafish valve morphogenesis through fibronectin synthesis. Nature Communications 2016 7:1 7, 1–14 (2016).

53. Kale, G. R. et al. Distinct contributions of tensile and shear stress on E-cadherin levels during morphogenesis. Nature Communications 2018 9:1 9, 1–16 (2018).

54. Baeyens, N., Bandyopadhyay, C., Coon, B. G., Yun, S. & Schwartz, M. A. Endothelial fluid shear stress sensing in vascular health and disease. J Clin Invest 126, 821–828 (2016).

55. Lee, H. J. et al. Fluid shear stress activates YAP1 to promote cancer cell motility. Nature Communications 2017 8:1 8, 1–14 (2017).

56. Lachowski, D. et al. FAK controls the mechanical activation of YAP, a transcriptional regulator required for durotaxis. The FASEB Journal 32, 1099–1107 (2018).

57. Lin, D. C., Dimitriadis, E. K. & Horkay, F. Robust Strategies for Automated AFM Force Curve Analysis—I. Non-adhesive Indentation of Soft, Inhomogeneous Materials. J Biomech Eng 129, 430–440 (2007).

58. Guizar-Sicairos, M., Thurman, S. T. & Fienup, J. R. Efficient subpixel image registration algorithms. Opt Lett 33, 156–158 (2008).

## REFERENCE

[1] S. Yoshida, ‘Waves: Fundamentals and dynamics’, Waves: Fundamentals and dynamics, 2017, doi: 10.1088/978-1-6817-4573-2.

[2] J. M. Walker et al., ‘Nondestructive evaluation of hydrogel mechanical properties using ultrasound’, Ann Biomed Eng, vol. 39, no. 10, pp. 2521–2530, Oct. 2011, doi: 10.1007/S10439-011-0351-0/FIGURES/7.

[3] A. Ahmad et al., ‘Magnetomotive optical coherence elastography using magnetic particles to induce mechanical waves’, Biomedical Optics Express, Vol. 5, Issue 7, pp. 2349-2361, vol. 5, no. 7, pp. 2349–2361, Jul. 2014, doi: 10.1364/BOE.5.002349.

[4] J. E. Sader, ‘Frequency response of cantilever beams immersed in viscous fluids with applications to the atomic force microscope’, J Appl Phys, vol. 84, no. 1, pp. 64–76, Jul. 1998, doi: 10.1063/1.368002.

[5] W. Tyrrell. Thomson and M. Dillon. Dahleh, ‘Theory of vibrations with applications’, p. 524, 2014.

[6] G. R. Cowper, ‘The Shear Coefficient in Timoshenko’s Beam Theory’, J Appl Mech, vol. 33, no. 2, pp. 335–340, Jun. 1966, doi: 10.1115/1.3625046.

[7] V. Giurgiutiu, ‘Structural health monitoring with piezoelectric wafer active sensors’, 16th International Conference on Adaptive Structures and Technologies, pp. 94–100, 2006, doi: 10.1016/C2013-0-00155-7.

[8] B. A. (1922-). Auld, ‘Acoustic fields and waves in solids. Vol. 2’, 1990, Accessed: Jul. 17, 2023. [Online]. Available: https://books.google.com/books/about/Acoustic_Fields_and_Waves_in_Solids.html?id=qUGEPwAACAAJ

[9] C. E. Repetto, A. Roatta, and R. J. Welti, ‘Forced vibrations of a cantilever beam’, Eur J Phys, vol. 33, no. 5, p. 1187, Jul. 2012, doi: 10.1088/0143-0807/33/5/1187.

[10] H. Tao et al., ‘Oscillatory cortical forces promote three dimensional cell intercalations that shape the murine mandibular arch’, Nat Commun, 2019, doi: 10.1038/s41467-019-09540-z.

[11] E. H. Barriga, K. Franze, G. Charras, and R. Mayor, ‘Tissue stiffening coordinates morphogenesis by triggering collective cell migration in vivo’, Nature, vol. 554, no. 7693, pp. 523–527, 2018, doi: 10.1038/nature25742.

[12] M. Zhu et al., ‘Spatial mapping of tissue properties in vivo reveals a 3D stiffness gradient in the mouse limb bud’, Proc Natl Acad Sci U S A, 2020, doi: 10.1073/pnas.1912656117.

[13] K. Bambardekar, R. Clément, O. Blanc, C. Chardès, and P.-F. Lenne, ‘Direct laser manipulation reveals the mechanics of cell contacts in vivo’, Proceedings of the National Academy of Sciences, 2015, doi: 10.1073/pnas.1418732112.

[14] G. Scarcelli et al., ‘Noncontact three-dimensional mapping of intracellular hydromechanical properties by Brillouin microscopy’, Nat Methods, 2015, doi: 10.1038/nmeth.3616.

[15] S. Möllmert et al., ‘Beyond comparison: Brillouin microscopy and AFM-based indentation reveal divergent insights into the mechanical profile of the murine retina’, bioRxiv, p. 2024.01.24.577013, Jan. 2024, doi: 10.1101/2024.01.24.577013.

[16] K. M. Kennedy et al., ‘Quantitative micro-elastography: imaging of tissue elasticity using compression optical coherence elastography’, Scientific Reports 2015 5:1, vol. 5, no. 1, pp. 1–12, Oct. 2015, doi: 10.1038/srep15538.

[17] V. Y. Zaitsev et al., ‘Strain and elasticity imaging in compression optical coherence elastography: The two-decade perspective and recent advances’, J Biophotonics, vol. 14, no. 2, p. e202000257, Feb. 2021, doi: 10.1002/JBIO.202000257.

[18] A. Tang, G. Cloutier, N. M. Szeverenyi, and C. B. Sirlin, ‘Ultrasound elastography and MR elastography for assessing liver fibrosis: Part 1, principles and techniques’, American Journal of Roentgenology, vol. 205, no. 1, pp. 22–32, Jul. 2015, doi: 10.2214/AJR.15.14552/SUPPL_FILE/07_15_14552_SUPPDATA.PDF.

[19] R. Loomba et al., ‘Novel 3D Magnetic Resonance Elastography for the Noninvasive Diagnosis of Advanced Fibrosis in NAFLD: A Prospective Study’, American Journal of Gastroenterology, vol. 111, no. 7, pp. 986–994, Jul. 2016, doi: 10.1038/AJG.2016.65.

